# Novel DNA endonuclease activity of human ERK1 protein

**DOI:** 10.1101/2025.03.03.641133

**Authors:** Heeyoun Bunch, Sangmin Ju, Jaehyeon Jeong, Bin Chan Joo, Min Ji Yu, Dahun Lee, Ki-Soon Yoon, Sanzhar Tarassov, Jeong Ho Chang, Jae-Han Jeon, Stuart K. Calderwood, Yeon Ui Lee

## Abstract

ERK1 is a key kinase in the mitogen-activated protein kinase pathway, which transduces extracellular signals into cellular responses. Following triggering of the pathway, cytosolic ERK1 is phosphorylated, translocates into the nucleus, and activates DNA-bound transcription factors and target genes. This study reports a novel, non-canonical role of human ERK1 as a DNA endonuclease. It catalyzes single-strand DNA breaks in sequence-unspecific yet, topology-sensitive manners. ERK1 catalysis requires magnesium ions and no ATP consumption. Comprehensive biochemical and *in silico* structural analyses indicate that a Mg^2+^ pocket, including a residue N^171^, which is allosterically modulated by the N-terminal domain, is important for DNA substrate binding and catalysis. Fluorescence imaging analysis suggests that ERK1 may interact with cytoplasmic DNAs in molecular condensates and destabilize them. Collectively, this study reveals an unprecedented role of ERK1 in nicking DNA molecules, which highlights the potential to regulate the fate of cytoplasmic DNAs.

## Introduction

The mitogen-activated protein kinase (MAPK) pathway consists of protein phosphorylation cascades that link extracellular signals to cellular responses, including gene regulation in plants and mammals (*1*). The MAPK pathway is crucial for controlling cell proliferation, DNA repair, apoptosis, and inflammation (*2*). Deregulation of the MAPK pathway is closely associated with cancer and neurodegenerative diseases; many compounds inhibiting protein kinases in the MAPK pathway have been developed and clinically tested for treatment of these pathologies (*3, 4*). Additionally, viral infections take advantage of MAPK pathway activation to promote the replication of the viral genome (*5*), indicating an important role of MAPK in viral and immune responses. MAPK1 and MAPK3 (ERK2 and ERK1, respectively) are phosphorylated and activated by MAPKK (MEK1/2) in the cytoplasm and are translocated into the nucleus, where they phosphorylate and activate nuclear transcription factors such as ELK1 and MYC (*3, 6–8*).

Our previous study revealed that ERK1 and ERK2 have different roles in the transcription of *EGR1* and *FOS*, representative immediate early genes (IEGs) and the first responder genes for cell cycle progression (*9*). ERK2 activates IEG transcription, whereas ERK1 represses it. Mechanistically, ERK2 stimulates topoisomerase IIβ (TOP2B) catalytic activity to relax positive DNA supercoiling and delays TOP2B to catalyze negative DNA supercoiling (*9*). Meanwhile, this study led us to speculate that ERK1 may possess a non-canonical enzymatic activity to resolve DNA supercoiling by itself, a hypothesis urgently and importantly needing validations. Thus, our present study has investigated the potential role of ERK1 as a DNA topology factor and nuclease, and further explored the physiological significance and implication of the new ERK1 function.

## Results

### Human ERK1 possesses DNA endonuclease activity

We compared first constitutively kinase-active human ERK mutants (*10*) —ERK1m (R^84^S) and ERK2m (R^67^S) — to investigate the basis for the differential properties of these enzymes in terms of endonuclease activities (*9*). These proteins were therefore expressed in *Escherichia coli* and purified using Ni affinity and gel filtration columns side-by-side (**fig. S1A**). Interestingly, ERK1m, but not the control (CTRL, ERK storage buffer only) or ERK2m, converted the negatively supercoiled substrate pBR322 (4361 bp) to nicked (or open-circular, OC) and linearized pBR322 plasmid in DNA relaxation assays (**Fig. 1A**). It should be noted that proteins were extracted by phenol, chloroform, and isoamyl alcohol immediately after the reaction, and so only DNA was separated and visualized on the gels in the DNA relaxation assay. To determine whether WT ERK1 also possessed a similar topological/DNase activity to that of ERK1m, ERK1 (K1) and ERK1m (K1m) were purified and compared side-by-side and confirmed by immunoblotting (**Fig. 1B**). TOP2B was included as a control to compare with ERK1s for their band patterns, as TOP2B generates partially relaxed plasmid species by removing DNA writhes one at a time (**fig. S1B**). The relaxation assay results showed that the enzymatic activities of ERK1 and ERK1m were comparable, yet distinctive from those of TOP2B (**Fig. 1, C** and **D**). ERK1 and ERK1m did not relax DNA supercoiling partially, as shown in the TOP2B reaction, but generated primarily nicked plasmids, then accumulated linearized ones (**Fig. 1C**). ERK1 did not require ATP to convert the supercoiled, circular plasmids to nicked or linear ones (**Fig. 1E**). In addition, the presence of magnesium ion (Mg^2+^) in reaction was critical for this endonuclease function of ERK1 and was not substitutable with other divalent ions, such as zinc or calcium (**Fig. 1F**).

**Fig. 1.**
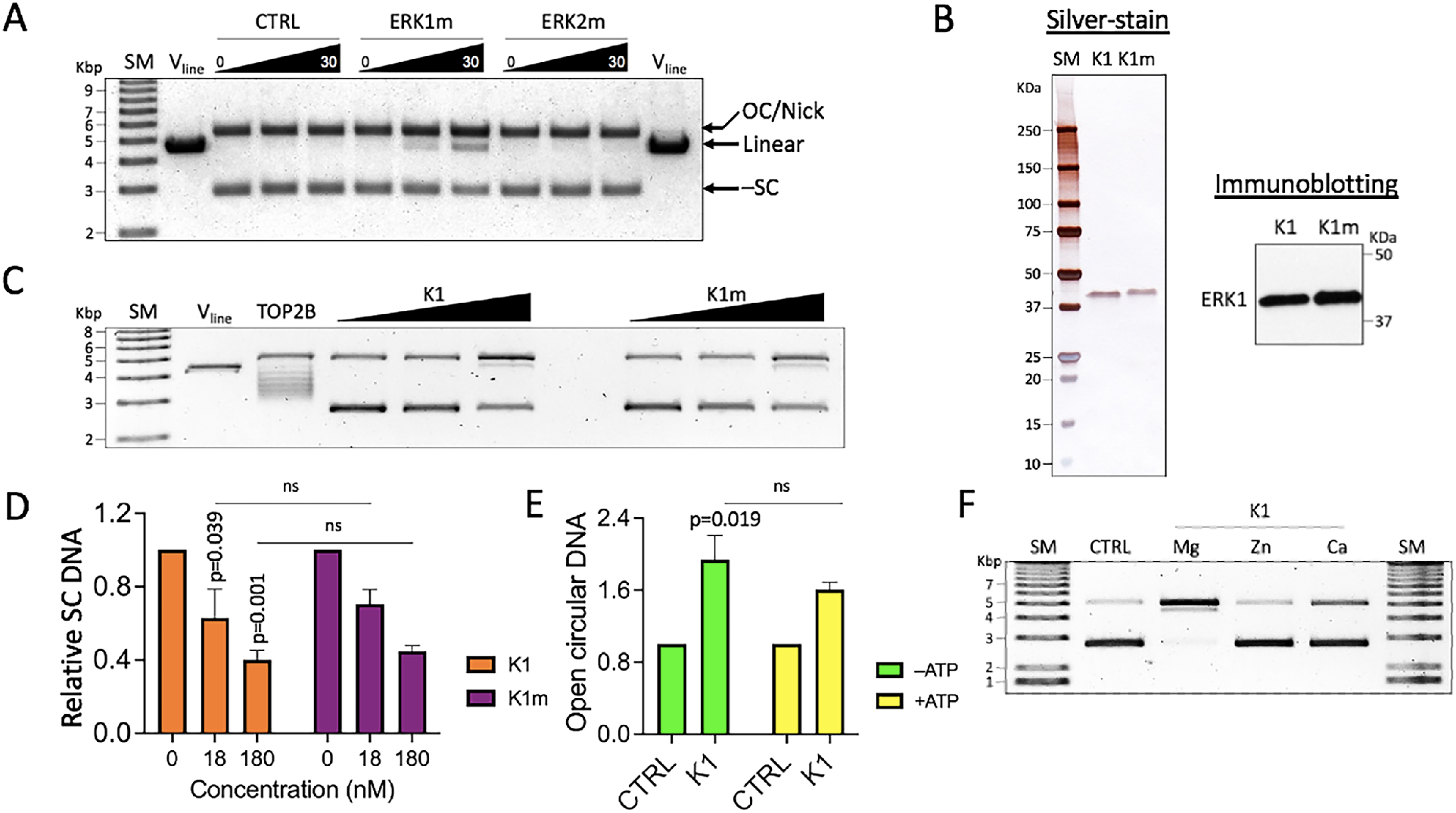
ERK1 relaxes circular, supercoiled DNA. (**A**) ERK proteins (200 ng or 4.6 pmol per each reaction) with a constitutively active kinase function (R^84^S [ERK1m] and R^67^S [ERK2m], respectively) were compared for their activities to relax pBR322 in time course, 0, 10, and 30 min. Unless otherwise indicated, 250 ng (3.53 nM) of pBR322 was used per a reaction. SM, size marker; V_line_, pBR322 linearized by HindIII; CTRL, ERK1 storage buffer only control; OC/nick, open-circular or nicked plasmid DNA; Linear, linearized plasmid DNA; –SC, negatively supercoiled plasmid DNA. (**B**) Purified ERK1 (K1) and ERK1m (K1m) proteins were visualized by silver-staining and their identity confirmed by immunoblotting. (**C**) Relaxation assay-gel showing both K1 and K1m comparably relax supercoiled pBR322 in a dose-dependent manner (0, 18, and 180 nM). pBR322 relaxed by TOP2B (2.8 fmol) was included as a comparison. (**D**) Quantification of relaxation assay results with K1 (orange bars) and K1m (purple bars) as mean values (n = 3) and standard deviation (SD). *P*-values were calculated with the unpaired, one sided Student’s t-test. ns, non-significant. (**E**) Relaxation assay data with K1 and pBR322 in reactions ± ATP (0 or 1 mM final concentrations, green and yellow bars, respectively). The data were presented as mean values (n = 3) and SD. *P*-values were calculated with the unpaired, one sided Student’s t-test. (**F**) Relaxation assay data showing K1 dependence on Mg^2+^ (Mg, 10 mM) for endonuclease activities. Zn^2+^ and Ca^2+^ were supplemented as 200 μM and 100 μM, respectively. Divalent ion final concentrations were determined considering their physiological concentrations. Note that linearized pBR322 DNA runs close to open circular one in (**C**) and (**F**) for the percentage of agarose gels (0.8%), compared to other gels (1.0 %).

### ERK1 catalyzes single-strand DNA breaks on DNA substrates with structural bias

To further characterize the DNA relaxing function, ERK1 (0, 2.5, 25, 50, 125, and 250 nM) and negatively supercoiled pBR322 were incubated for 3 h at 30 °C in DNA relaxation assays. Nicked and linearized plasmids, the catalyzed products, were accumulated in a dose-dependent manner (**Fig. 2, A** and **B** and **fig. S2A**). Nicked plasmid DNA was formed first, and then a portion of the products was converted to linearized ones (see 50, 125, and 250 nM in **Fig. 2A**). Time-course experiments with 230 nM ERK1 and 3.5 nM pBR322 plasmids showed almost complete supercoiled plasmid removal at 1 h timepoint and slowly increased levels of linearized bands at 3 and 5 h time points (**Fig. 2, C** and **D** and **fig. S2B**). Under the assumption that the substrates are circular, supercoiled DNA and that the products are the lost substrates converted to either nicked or linearized ones, K_cat_ of ERK1 is approximately 0.0005 min^−1^.

**Fig. 2.**
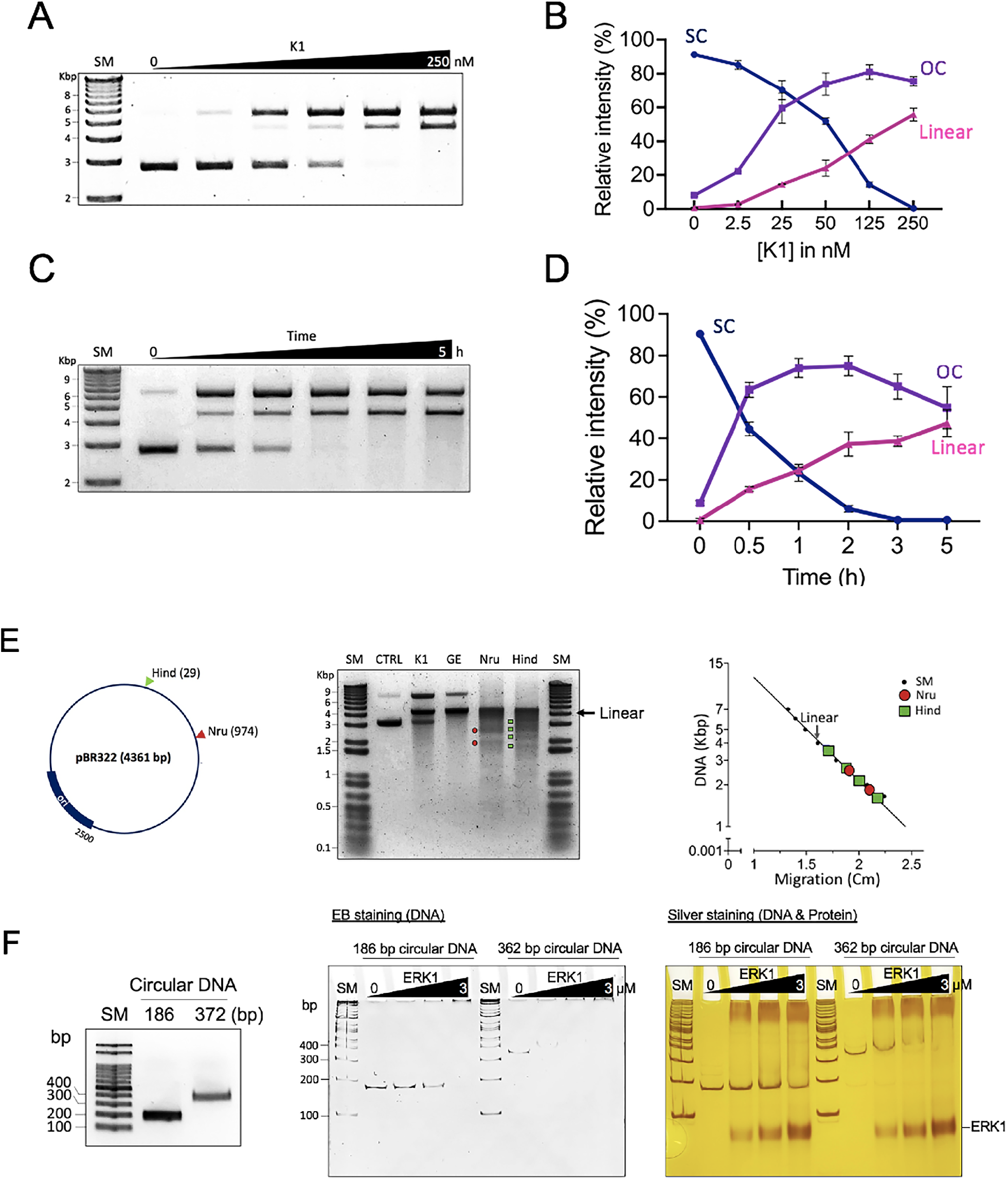
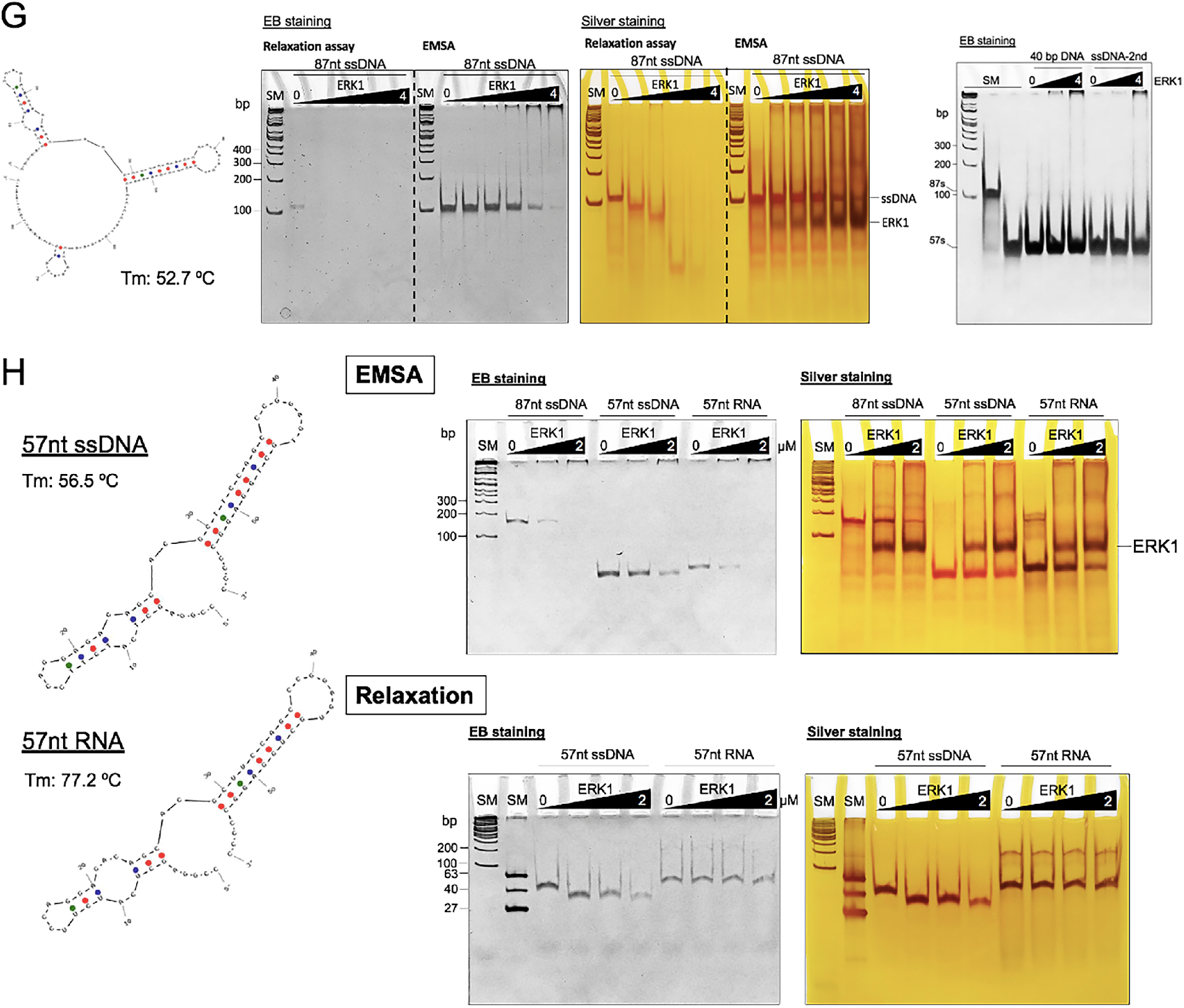
ERK1 catalyzes single-strand DNA break in DNA topology-specific manners. (**A**) Relaxation assay-gel showing ERK1 (K1) concentration titration (0, 2.5, 25, 50, 125, and 250 nM) for 3 h. (**B**) Quantification of the relaxation assay data of K1 concentration titration (n ≥ 3). Relaxation assay-gel showing pBR322 relaxation by K1 in time course analysis (0, 0.5, 1, 2, 3, 5 h). (**D**) Quantification of the relaxation assay data with K1 and pBR322 in time course analysis (n ≥ 3). (**E**) Left, pBR322 map showing HindIII (Hind), NruI (Nru), and *ori* sites. Middle, K1-mediated linearized pBR322 (K1) was gel-extracted (GE), and digested by NruI (Nru) or HindIII (Hind). CTRL, pBR322. Right, A linear regression line and equation derived from the relation between DNA migration distance and molecular weights of size marker DNA fragments (Kbp). The visible bands, after each restriction enzyme cut the K1-mediated linearized pBR322, were calculated based on the equation. (**F**) Left, small circular DNAs (186 and 372 bp) generated in this study. Middle, EMSA on a polyacrylamide gel stained with ethidium bromide (EB) showing small circular DNAs. WT ERK1 was titrated at 0, 0.75, 1.5, and 3 μM. Right, the same gel presented in the middle, subsequently stained by silver nitrate, showing both DNA and protein. (**G**) Left, the secondary structure of an 87nt-ssDNA used in this study. Middle (an identical gel in two staining methods), DNA relaxation assay and EMSA results, visualizing ssDNA digestion by ERK1 and ssDNA-ERK1 interaction stained by EB and silver nitrate. ERK1 was titrated at 0, 0.25, 0.5, 1, 2, and 4 μM concentrations and the DNA relaxation reaction was allowed for 3 h. Right, EMSA results with short, 40 bp dsDNA and ssDNA without specific secondary structures (ssDNA-2nd) and ERK1. ERK1 was titrated at 0, 2 and 4 μM concentrations. (**H**) A ssDNA and an RNA species with identical length and sequences (structures shown in the left) were compared for ERK1 binding (EMSA, top panels) and catalysis (relaxation assay, bottom panels). Polyacrylamide gels were stained by EB and then silver-nitrate. ERK1 was titrated at 0, 1, and 2 μM for EMSA and 0, 0.5, 1, and 2 μM for relaxation assays. The catalytic reaction was allowed for 2 h.

Next, we mapped the DNA sequences recognized for digestion by ERK1, employing Two enzymes, which cut pBR322 at only one restriction site, Hind III and Nru I (**Fig. 2E** and **fig. S2C**). We rationalized that a specific ERK1-catalyzed site(s) could be mapped if ERK1 linearized the circular plasmids in a sequence-specific manner, by cutting the ERK1-linearized DNA with these restriction enzymes. As shown in **Fig. 2E**, after pBR322 was incubated with ERK1 (lane 3; K1), the linearized plasmids were gel-purified (lane 4; GE) and digested by Hind III or Nru I. Either enzyme digestion produced multiple bands, accompanied by intensified smears rather than specific few bands, (lanes 5,6; Nru, Hind). These data suggested that ERK1 digests the plasmid at random sites in circular DNA in a sequence-unspecific manner. We speculate that random DNA breaking near the replication origin which is relatively underwound could produce the marked notable bands (e.g. 2536 and 1824 bp for Nru I) and faster-migrating DNA fragments with lower molecular weights below the linearized vector (**fig. S2D**).

Furthermore, we asked whether ERK1 possesses DNA conformation/topology-based substrate specificity as ERK1 displayed faster kinetics for supercoiled and open circular DNAs than linear ones in the relaxation assays. Small circular and linear double-stranded DNAs (dsDNAs) in different sizes, and single-stranded DNA (ssDNA) species with diverse lengths (27nt – 87nt) and secondary structure stabilities (30 ℃ < Tm < 60 ℃) were compared for ERK1 binding and catalysis. Circular DNAs sizing 186 bp and 372 bp were generated by ligating a 186 bp linear DNA with the *SacII* restriction enzyme site at its both ends (**Fig. 2F** and **fig. S2E**). As expected from pBR322-ERK1 interaction, ERK1 bound to these circular dsDNAs and linearized/digested them (**Fig. 2F** and **fig. S2F**). Strikingly, ERK1 bound to and fragmented all tested ssDNAs with different kinetics and digestion patterns. ERK1 rarely bound or digested short, linear dsDNAs below 40 bp (**Fig. 2G** and **fig. S2G**). We compared ERK1 interaction with a structured ssDNA vs an RNA chain with the identical sequence and length (57mer). Intriguingly, ERK1 bound to both, however it catalyzes the ssDNA but not the RNA counterpart (**Fig. 2H**). Notably, the RNA bound to ERK1 with a high affinity. In contrast, it barely bound ERK2 (**fig. S2H**). Collectively, these data revealed that ERK1 recognizes DNA conformation and binds preferentially to circular dsDNA or ssDNAs with secondary structures to catalyze them.

### N-terminal domain of ERK1 determines nucleolytic efficiency

Human ERK1 and ERK2 are 82.6% identical in amino acid sequences with distinctive N-terminal domains (NTD)(*11*). We therefore examined ERK1 and ERK2 N-terminal sequences for likely critical residues and mutated two consecutive arginines, at 15 and 16 to single or double alanine mutations within the region uniquely present in ERK1 (11–27 a.a.; **Fig. 3A**). The three mutant ERK1 proteins, R^15^A (15A), R^16^A (16A), or R^15^A/R^16^A (Double) mutants were bacterially expressed and purified (**Fig. 3B**). WT and mutant ERK1 proteins at concentrations 0, 60, 120, and 200 nM were incubated with negatively supercoiled pBR322 for 15 min before resolving the DNA molecules on the gel. 15A showed slightly inferior activity, whereas 16A ERK1 showed a notably enhanced nuclease activity, compared with the WT. The double mutant showed a similar activity to the one of the WT (**Fig. 3, C** and **D** and **fig. S3A**). Additionally, three residues, negatively charged glutamate, E^13^, E^18^, and E^27^ were individually mutated to alanine (**Fig. 3E**). The mutants were purified (**Fig. 3F**) and compared with the WT in the same experimental conditions, except for protein concentrations, as described in **Fig. 3C**. E^13^A (13A) and E^27^A (27A) exhibited enhanced nuclease activities, whereas E^18^A (18A) showed similar digestion efficiencies, compared to the WT (**Fig. 3, G** and **H** and **fig. S3B**). These data suggested that the N-terminal region spanning residues 11–27 suppresses the nuclease activity of ERK1 involving electrostatic interactions through E^13^, R^16^, and E^27^.

**Fig. 3.**
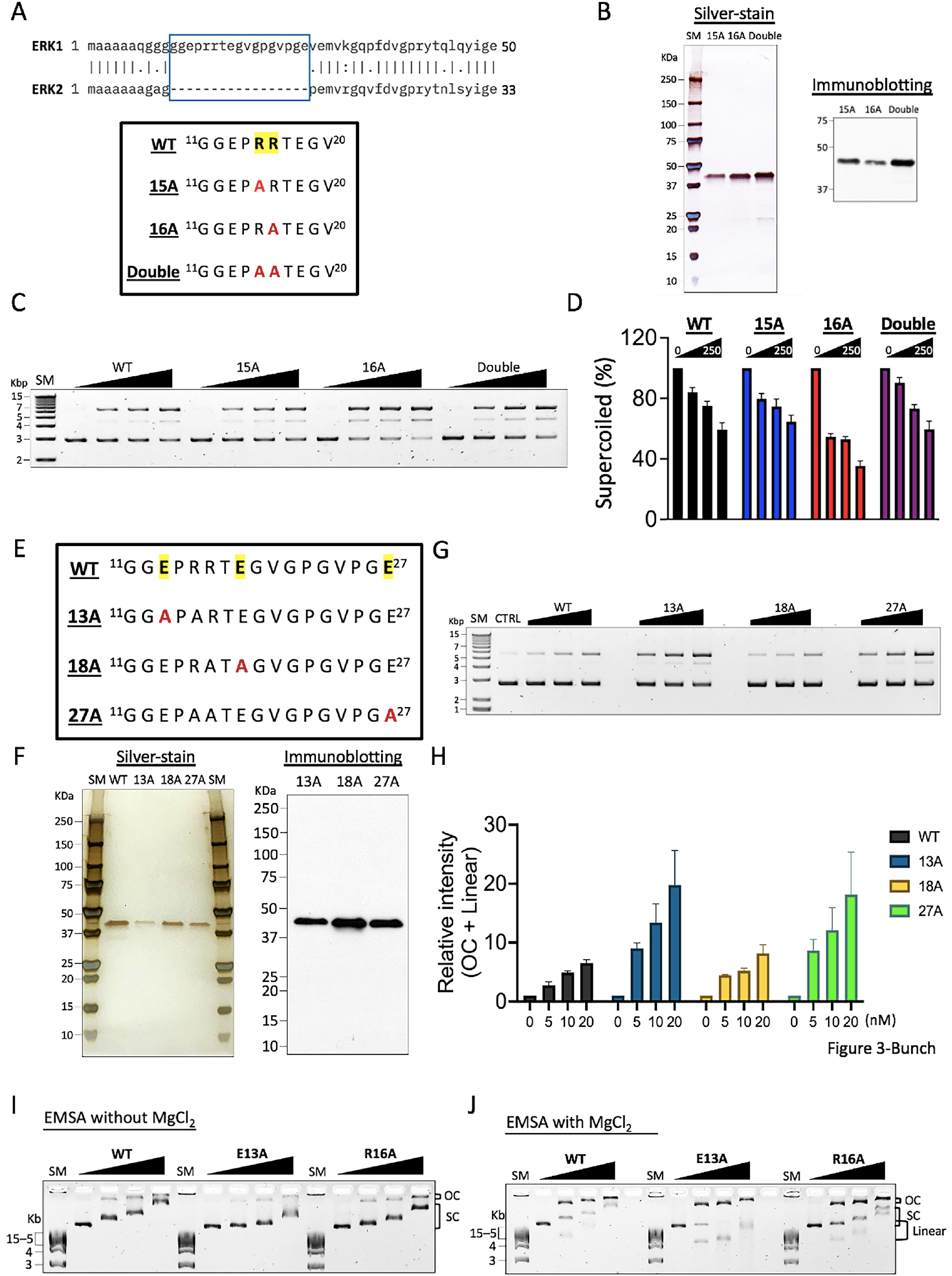
NTD of ERK1 determines nucleolytic efficiency. (**A**) Top, comparison of the NTD of ERK1 and ERK2. The region present only in ERK1 was boxed in sky-blue (11 – 27 a.a.). Bottom, R^15^ (R15) and R^16^ (R16) were marked with yellow. Three missense mutants with alanine substitutions were depicted as letter A (alanine) in red. (**B**) Purified R^15^A (15A), R^16^A (16A), and Double (R^15^A/R^16^A) mutant ERK1 proteins were visualized by silver-staining and confirmed by immunoblotting. (**C**) pBR322 relaxation by ERK1 mutants for 15 min in concentration titration (0, 60, 120, and 200 nM). (**D**) Quantification of reduced negatively supercoiled pBR322 bands by WT (black bars), 15A (blue bars), 16A (red bars), and Double mutant (purple bars) ERK1 in the concentration titration analyses (n = 3). (**E**) Three glutamates of E^13^, E^18^, and E^27^ (marked with yellow) were substituted to alanines (A, marked in red) (**F**) Purified E^13^A (13A), E^18^A (18A), and E^27^A (27A) ERK1 proteins were visualized by silver-staining and confirmed by immunoblotting. (**G**) Concentration titration of WT, 13A, 18A, and 27A (0, 5, 10, and 20 nM; 15 min reaction time). (**H**) Quantification of increased OC or linear pBR322 bands by WT (grey), 13A (blue), 18A (yellow), and 27A (green) ERK1 in the concentration titration analyses (n = 3). **(I)** EMSA with pBR322 (1 nM) and WT, 13A, and 16A ERK1 (0, 1, 2.5, and 5 μM) in the absence of MgCl_2_, and (**J**) in the presence of MgCl_2_.

Binding affinity to substrate, pBR322, was next monitored for WT, 13A, and 16A ERK1 proteins using the gel electrophoretic mobility shift assay (EMSA). Agarose gels, instead of polyacrylamide ones, were used for the assay owing to a large molecular weight (4361 bp) and bulky shapes of the plasmid DNA (*12*). The WT or mutant ERK1s were titrated at concentrations 0, 1, 2.5, and 5 μM with 1 nM pBR322 for 10 min at room temperature before loading onto the gel. Despite the large size and bulky shape of DNA, higher gel percentages were needed to visualize band-shift because of the molecular weight of ERK1 (44 KDa), which is equivalent to approximately 70 bp dsDNA. These challenges limited the resolution only to linearized DNAs with sizes below 5 Kbp (see SM in **Fig. 3, I** and **J**). In the absence of Mg^2+^, a dose-dependent band shift was observed, with major bands for the SC DNA-protein complexes and minor bands for the OC DNA-protein complexes (**Fig. 3I**). The two mutants, 13A and 16A, and especially 13A, appeared to have a reduced binding affinity to the SC DNA, compared to the WT (**Fig. 3I**). In the presence of Mg^2+^, SC, linear, and OC DNA bands were shown, as expected, because ERK1 could actively catalyze SC DNAs into OC and OC to linear DNAs (**Fig. 3J**). Despite showing a lower substrate binding affinity without Mg^2+^, 13A ERK1 produced linear DNA more efficiently than WT or 16A in the presence of Mg^2+^ (**Fig. 3J**). All three proteins bound to SC, OC, and linear plasmid DNAs to shift their mobility, yet with different affinities and catalytic efficiencies (**Fig. 3J**).

### Artificial intelligence-predicted ERK1 structure model

We investigated the ERK1 protein sequence and found that it lacks known motifs of DNA endonuclease or nickase. Additionally, currently available human ERK1 and ERK2 structures have limited information because they were co-crystalized with other peptides and/or inhibitors and omitted the apparently flexible N- and C-terminal domains (*13, 14*). We attempted but did not succeed to resolve the ERK1 structure through cryo-electron microscopy. Therefore, we employed an artificial intelligence (AI)-based tool, AlphaFold 3 to predict ERK protein structures and to evaluate the structural potentials to interact with and catalyze DNA molecules (*15*). AI-predicted full-length ERK1 (379 a.a., 44 KDa) and ERK2 (360 a.a., 42 KDa) structures showed overall similarities and yet distinctive NTDs and surface contours (**Fig. 4A** and **fig. S4A**). Next, ERK1 was co-structured with DNA segments of pBR322. It was found that ERK1-DNA interaction was established only in the inclusion of Mg^2+^ in the algorithm (**fig. S4B**). Interestingly, the structure of the ERK1-DNA (pBR322 DNA segments including 2292–2319)-Mg^2+^ complex indicated a DNA binding cleft and a Mg^2+^ binding site in ERK1, where N^171^ and D^184^ could have electrostatic interactions with Mg^2+^ within approximately 2.5 Å proximities (**Fig. 4B** and **fig. S4C**). Mutating N^171^ and D^184^ to alanine predictably dissociated Mg^2+^ from the binding pocket (**fig. S4D**). The distance between Mg^2+^ and the DNA backbone was approximately 6.8 Å, fluctuating, dependent on the input DNA segments (**Fig. 4B**). The predicted local distance difference test (pLDDT) of the ERK1-DNA complex indicated high model confidences (pLDDT > 70) for the middle domain, including the Mg^2+^ binding site, for example, N^171^ and D^184^, and the enzyme-DNA interface (**Fig. 4C**).

**Fig. 4.**
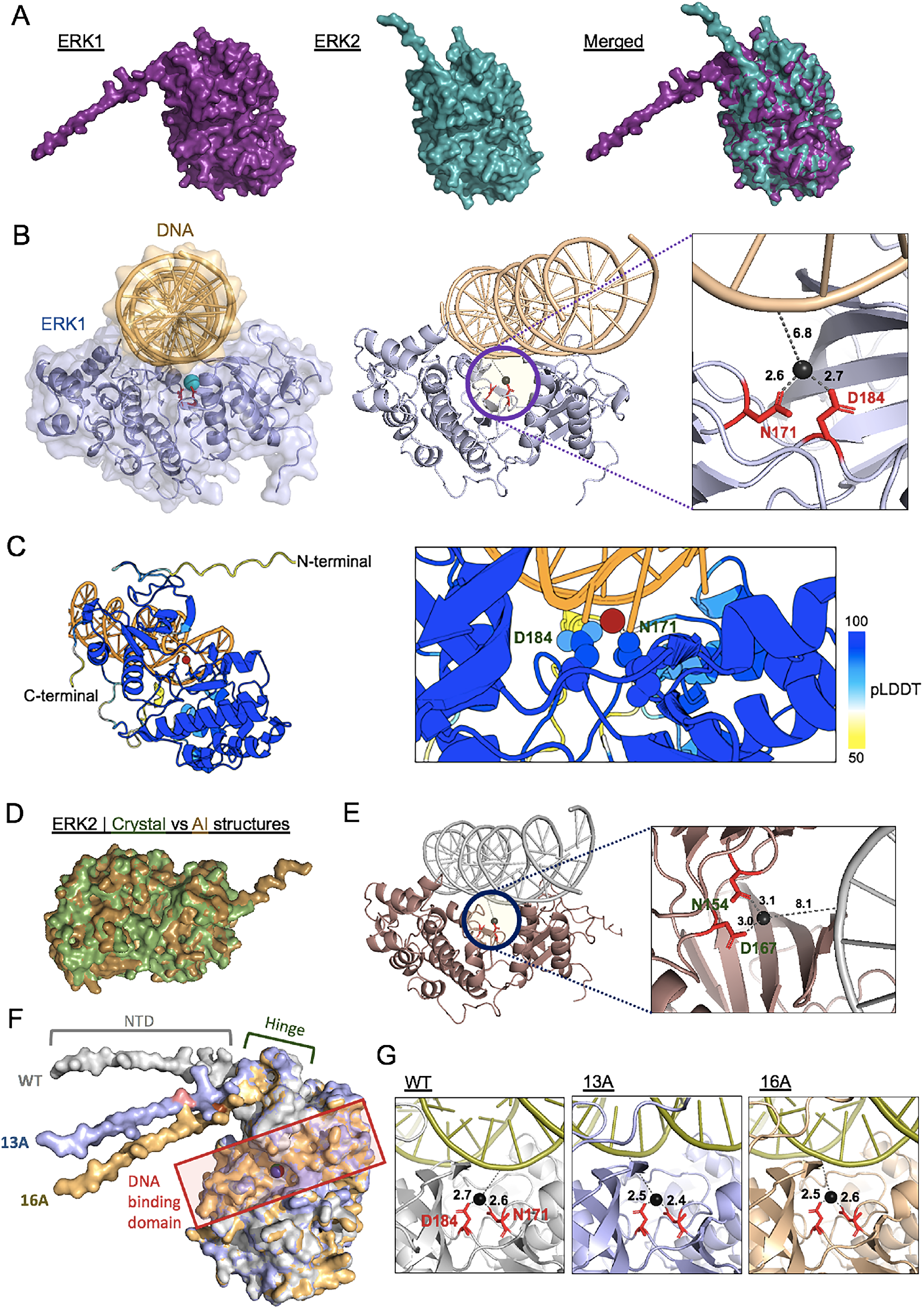
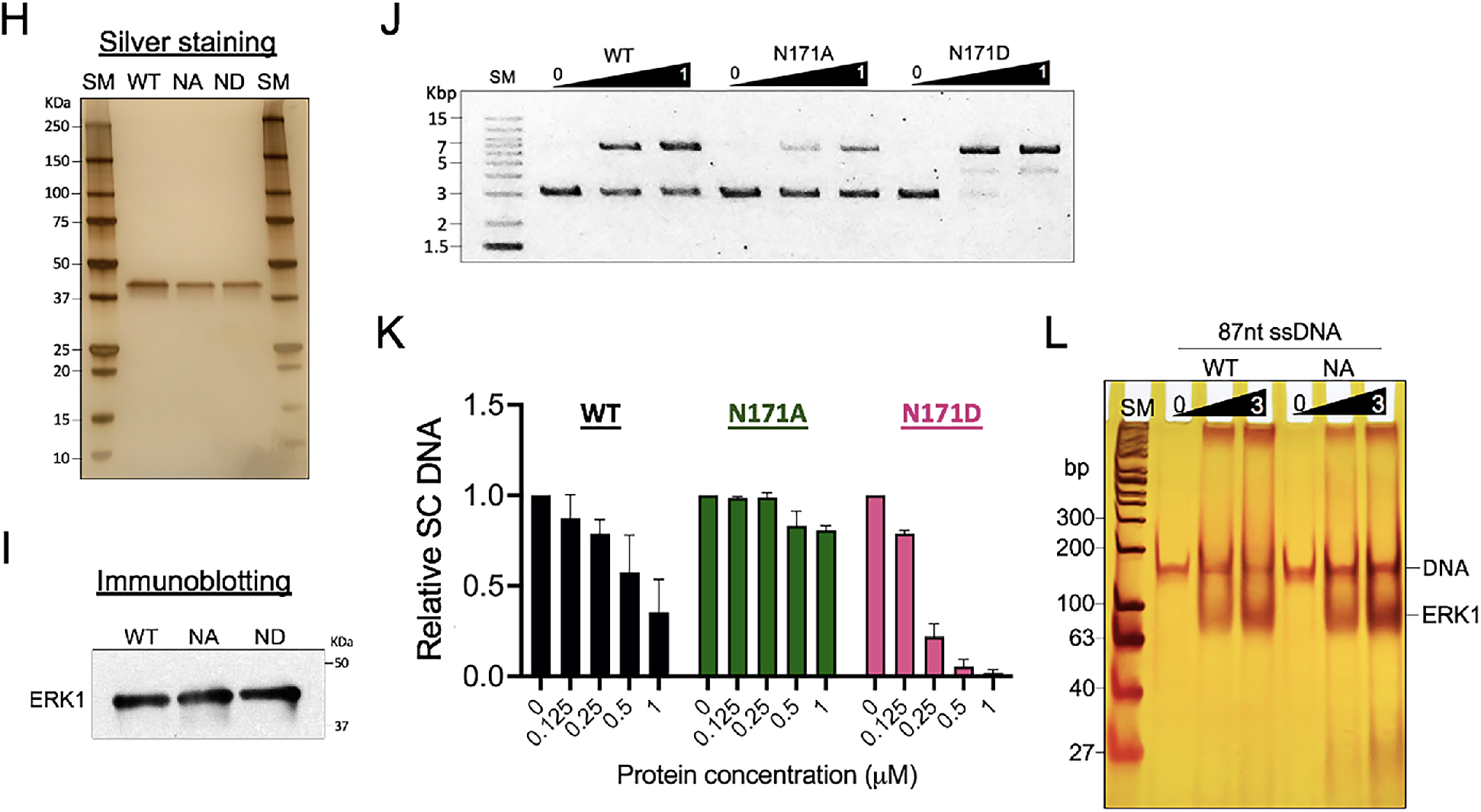
AI-predicted ERK1 structure model. (**A**) Left, AI-predicted ERK1 structure; middle, AI-predicted ERK2 structure; right, ERK1-ERK2 merged structures. The Cα root mean square deviation (RMSD) between ERK1 and ERK2, 0.504 Å. (**B**) Left, 3D structure of ERK1 (silver) and DNA (tan) complex in space-fills. Mg^2+^ in sky blue. N^171^ and D^184^ in red sticks. Middle, the complex omitting space-fills and the area with a purple circle showing the putative catalytic site of ERK1 as an endonuclease and in a zoom-in view (right). Mg^2+^, a black sphere. Atomic distances (Å) shown in gray, dashed lines and numbers. ModelArchive ID: ma-0se3b. (**C**) AI-predicted ERK1-DNA (87nt ssDNA) complex reflecting pLDDT scores (50 – 100). Left, Mg^2+^ in red, DNA in orange, N^154^ and D^167^ shown in sticks. The unstructured N-terminal region was partly truncated. Right, a zoom-in view of the Mg^2+^ and DNA binding site. N^171^ and D^184^ shown in space-fills. ModelArchive ID: ma-tav35. (**D**) Merged crystal (green, PDB ID: 4H3Q) and AI-predicted (gold) ERK2 structures. RMSD, 0.435 Å. (**E**) AI-predicted ERK2-DNA complex with a blue circle showing a Mg^2+^ (A black sphere) binding site (left) and a zoom-in view (right). N^154^ and D^167^ shown in red sticks. ModelArchive ID: ma-p6dyo. (**F**) Merged structures of WT (light gray), 13A (light blue), and 16A (light orange) ERK1. The E^13^A residue in 13A and the R^16^A residue in 16A were marked in pink and orange, respectively. Mg^2+^ in black (WT), dark blue (13A), and red (16A); DNA binding cleft/domain boxed in red. (**G**) Comparison of distances between Mg^2+^ and two coordinating amino acids, N^171^ and D^184^ in WT and 13A and 16A mutant ERK1 proteins. The same DNA (28-mer, 2292–2319 of pBR322) was used in **Fig. 4, B, E**, and **G**. (**H**) Purified WT, N^171^A (NA), and N^171^D (ND) ERK1 shown by silver-staining. (**I**) WT, NA, and ND ERK1 verified by immunoblotting. (**J**) Representative relaxation assay results showing notably reduced and increased DNA relaxation of pBR322 by NA and ND mutations, respectively. ERK1 was titrated to final concentrations of 0, 0.5, and 1 μM and the reaction was incubated for 30 min. (**K**) Bar graphs comparing WT, NA, and ND ERK1 for their effectiveness to relax the negatively supercoiled pBR322. (**L**) Representative EMSA result on a polyacrylamide gel stained with silver nitrate showing compromised DNA (87nt ssDNA) binding of NA ERK1, compared to the WT.

A comparison between the crystal (complexed with a docking peptide and missing N- and C-terminal domains, PDB ID: 4H3Q) and AI-predicted (full-length) structures of ERK2 showed overall similarities, supporting confidence in AI structures (**Fig. 4D** and **fig. S4E**). Similar to those of ERK1, AI-predicted ERK2 structures showed DNA (2292–2319 of pBR322) and Mg^2+^ binding sites (**Fig. 4E**). However, the distances between Mg^2+^ and N^154^ and D^167^ and between Mg^2+^ and DNA backbone were overall farther, 3.1 Å and 8.1 Å, respectively, compared to those of ERK1, suggesting weaker interactions (**Fig. 4E**). In addition, we compared AI-predicted WT, 13A, and 16A ERK1 structures, which revealed that these mutants rotated the flexible NTD in different angles, allosterically modulating the conformation of the hinge and DNA binding domains to shorten the distances between the critical Mg^2+^ ion and the two coordinating amino acids (**Fig. 4, F** and **G** and **fig. S4F**). Mechanistically, the structures suggested that the strength of Mg^2+^ binding to the ERK1 enzyme might be important for catalytic efficiency. Supporting the model, N^171^D (ND) mutation strengthened the nuclease activity of the enzyme, whereas N^171^A (NA) weakened the substrate binding and catalysis, compared to the WT ERK1 (**Fig. 4, H–L** and **fig. S4G**). We speculated the possibility of hydrated Mg^2+^ or titratable residues near the DNA binding site, such as Y^53^, K^168^, or S^170^, which were located within 5 Å from the Mg^2+^ ion, for the mediation of DNA strand break (**fig. S4, H** and **I**), which awaits further investigation.

### ERK1 interacts with and destabilizes cytoplasmic DNA

Our previous study showed that bacterially purified, recombinant ERK1 bound to the DNA *in vitro*, whereas nuclear ERKs from HeLa nuclei did not (**fig. S5A)**(*9*). One possible difference between these two protein groups is post-translational modification, as ERK1 is known to be translocated into the nucleus in a phosphorylation-dependent manner (*16*). Additionally, recent studies have identified a variety of cytoplasmic DNA (cytoDNA) species in humans, including extrachromosomal DNA, mitochondrial DNA, micronuclei, cytoplasmic chromatin fragments, and viral and bacterial DNA (*17–22*). Therefore, we hypothesized that ERK1 in the cytoplasm might interact with cytoDNAs. To test this hypothesis, ERK1 and cytoDNA were monitored in HEK293T cells under a high-resolution fluorescence microscopy using an ERK1 antibody and over-exposed DAPI staining (*23–26*), respectively. DNA-RNA probes generated based on chromosome 7p11.2 loci were tested as controls (**fig. S5B**), which confirmed DAPI staining specific to DNAs, but not RNAs. ERK1 was located primarily in the cytoplasm, and a portion of the ERK1 proteins were colocalized with dsDNAs (F1–3, **Fig. 5A** and **fig. S5C**). Super-resolution imaging of focal spots, F1–3, strikingly, showed that a large number of ERK1 molecules (44 KDa in approximately 2.4 nm) interacted with thousands of dsDNA molecules, each sizing approximately 20 nm, which is equivalent to approximately 55 Kbp dsDNA, forming nano-scale condensates (**Fig. 5B**).

**Fig. 5.**
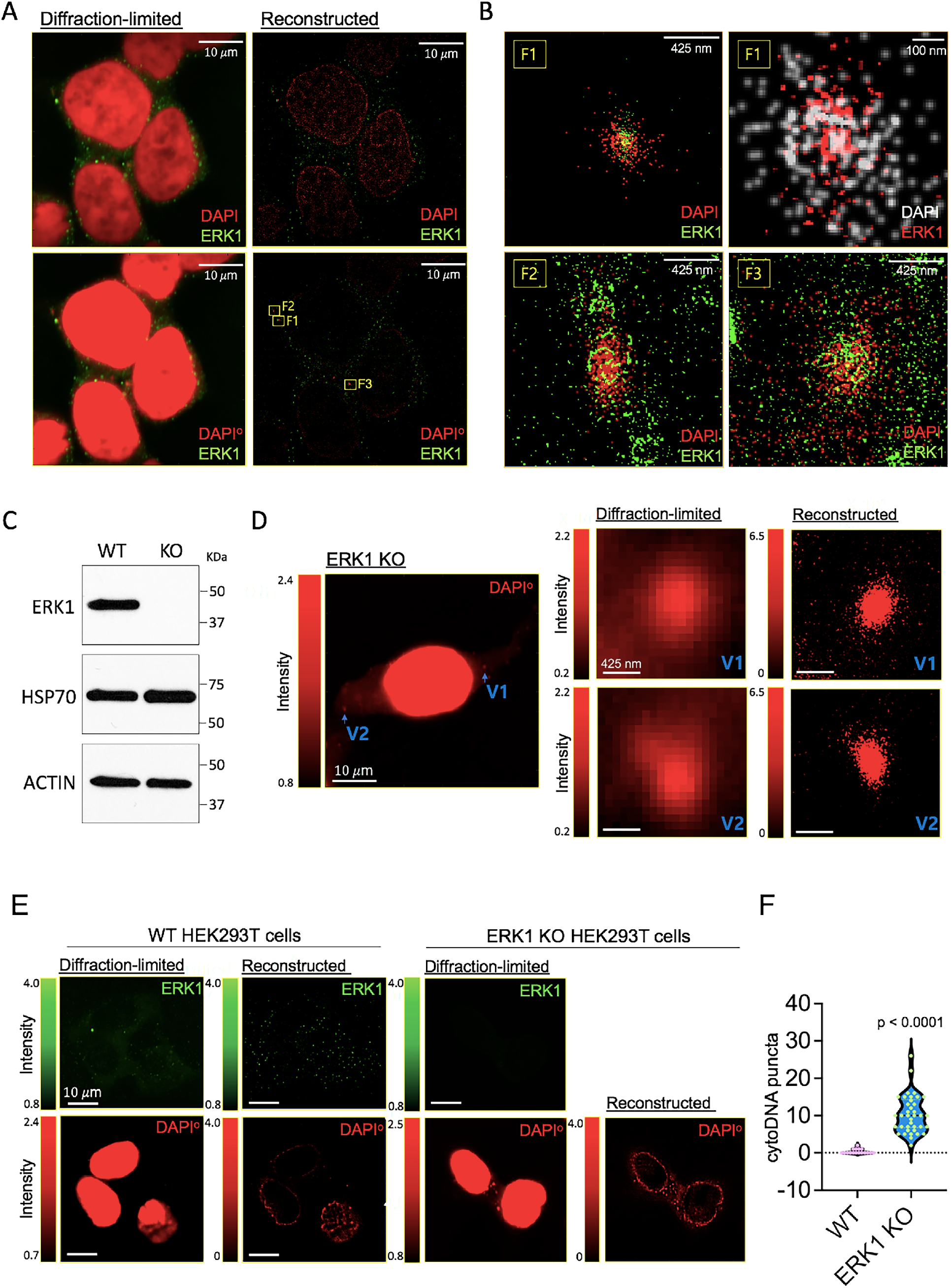
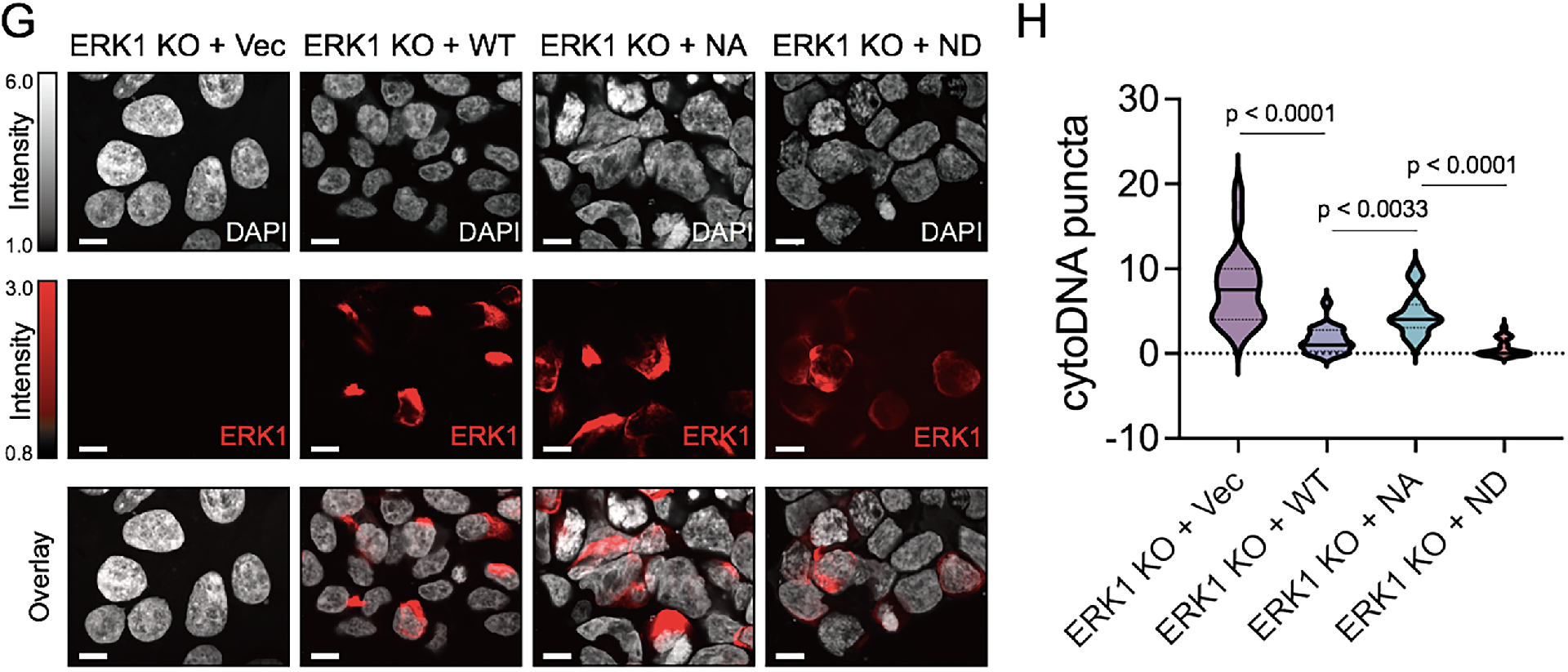
ERK1 interacts with and destabilizes cytoplasmic DNA. (**A**) Diffraction-limited and reconstructed super-resolution multicolor images of HEK293T cells. Top, diffraction-limited (left) and super-resolution (right) fluorescence images. DAPI (red) and ERK1 (Alexa 647, green). Bottom, overexposed DAPI (DAPI°) merged with ERK1 (left) and the super-resolution image excluding oversaturated DAPI signals (right). F1–3, focal sites of DAPI and ERK1 overlapping signals. Intensity scales should be multiplied by 10^4^ in arbitrary units (AU) throughout **Fig. 5**. (**B**) Zoom-in views of F1–3 showing cytoDNA-ERK1 interaction. (**C**) Validation of WT and ERK1 KO HEK293T cells by immunoblotting. β-ACTIN and HSP70 as reference and loading controls. (**D**) Cytoplasmic DAPI puncta (red) in an ERK1 KO HEK293T cell (left). V1–2 sites were magnified in diffraction-limited and super-resolution microscopy images (right). (**E**) Diffraction-limited and super-resolution microscopy images of WT and ERK1 KO cells showing colocalization of ERK1 (Alexa 647, green) and DNA (DAPI°, red). (**F**) Quantification of cytoDNA puncta revealing a significant increase in ERK1 KO cells (blue violin, n = 38) compared to WT cells (pink violin, n = 17). The data were presented as mean values and SD. *P*-values were calculated with the unpaired, two-sided Student’s t-test. (**G**) WT, NA, and ND ERK1 proteins tagged with GFP were reintroduced into ERK1 KO cells: ERK1 KO + WT, NA, and ND, respectively. DAPI, gray; ERK1, red; ERK1 KO + Vec, ERK1 KO HEK293T cells transfected with a GFP vector control. A scale bar, 5 μm. (**H**) CytoDNA puncta were counted in a GFP only control (KO + Vec, n = 20) and WT (ERK1 KO + WT, n = 20), NA (ERK1 KO + NA, n = 20), and ND (ERK1 KO + ND, n = 20) ERK1-expressing ERK1 KO HEK293T cells. *P*-values were calculated with two-way analysis of variance (ANOVA) with Tukey’s multiple comparisons test. Medians, black solid lines; quartiles, dotted black lines.

Next, based on the evidence that ERK1 digests and destabilizes circular and linear DNAs with its DNA endonuclease activity *in vitro*, we hypothesized and evaluated whether ERK1 knock-out (KO) increases the amount of cytoDNAs and their puncta in cells. The analysis using CRISPR-Cas 9-edited ERK1 KO HEK293T cells showed that cytoDNA puncta, which consisted of a number of dsDNA molecules, were more readily visible and significantly increased in ERK1 KO cells, compared to the WT HEK293T cells (**Fig. 5, C–F** and **fig. S5D**). Importantly, reintroducing WT ERK1 in ERK1 KO cells notably reduced cytoDNA puncta counts (**Fig. 5, G** and **H** and **fig. S5E**). Compared to WT ERK1, NA ERK1, which has a compromised nuclease function *in vitro* (**Fig. 4, J–L**), showed more visible cytoDNA puncta, whereas these were markedly diminished in the KO cells replenished with ND ERK1 (**Fig. 5, G** and **H**).

## Discussion

Collectively, these findings support a novel, non-canonical enzymatic function of human ERK1 protein as a DNA endonuclease that nicks and digests dsDNA and structured ssDNA. ERK1 is capable of mediating dsDNA break and destabilization through single stranded DNA breakage. Notably, the catalysis of protein kinases involves hydrolyzing ATP to transfer the γ-phosphate of the purine nucleotide triphosphate to the hydroxyl group of their substrates covalently (*27*). The endonuclease function of ERK1, a critical transcription factor and kinase, raises a question: whether more DNA-binding transcription factors with phosphotransferase function might also be able to break the DNA strand by such redox reactions. Our AI-based structural model illustrates the ERK1-DNA interaction and the important role of Mg^2+^ for the ERK1-mediated DNA catalysis. DNA break is a critical cellular event, linked to DNA topological changes and recombination, gene regulation, genomic mutation/instability, microbial defense, cell fate, and more (*28–33*). Therefore, our finding that ERK1 has a novel endonuclease activity is important and likely associates with a variety of physiological and clinical implications. Finally, cytoDNAs have emerged as critical regulators of inflammation, immunity, and degenerative diseases (*34–36*). ERK1-cytoDNAs interaction shown in this study suggests potentially important roles of cytoplasmic ERK1 in regulating the fate and topology of cytoDNAs, which awaits future studies.

## Acknowledgments

We thank D. Engelberg and N. Soudah at the Hebrew University of Jerusalem in Israel for their generosity in sharing bacterial WT and mutant ERK1 and ERK2 expression vectors. We appreciate D.J. Taatjes at the University of Colorado in the USA and S. Sekine at RIKEN in Japan for critical reading and discussions and J. Jung, J. Yoo, and former and current members of the Bunch laboratory at Kyungpook National University (KNU) for their technical assistance. H.B thanks J. Bunch and J. Christ for their loving encouragement and support throughout the course of this work.

## Author contributions

Conceptualization: HB

Methodology: JHC, YUL, HB

Investigation: SJ, JJ, BJ, MY, KY, DL, ST, HB

Visualization: SJ, JJ, HB

Funding acquisition: J-HJ, HB

Project administration: J-HJ, YUL, SKC, HB

Supervision: JHC, J-HJ, YUL, SKC, HB

Writing – original draft: HB

Writing – review & editing: YUL, SKC, HB

## Data and materials availability

All data are available in the main text or the supplementary materials.

